# Regulation of Microtubule Disassembly by Spatially Heterogeneous Patterns of Acetylation

**DOI:** 10.1101/725895

**Authors:** J S Aparna, Ranjith Padinhateeri, Dibyendu Das

**Affiliations:** Centre for Research in Nanotechnology and Science, Indian Institute of Technology Bombay, Mumbai, India; Department of Biosciences and Bioengineering, Indian Institute of Technology Bombay, Mumbai, India; Department of Physics, Indian Institute of Technology Bombay, Mumbai, India 400076

## Abstract

Microtubules (MTs) are bio-polymers, composed of tubulin proteins, involved in several functions such as cell division, transport of cargoes within cells, maintaining cellular structures etc. Their kinetics are often affected by chemical modifications on the filament known as Post Translational Modifications (PTMs). Acetylation is a PTM which occurs on the luminal surface of the MT lattice and has been observed to reduce the lateral interaction between tubulins on adjacent protofilaments. Depending on the properties of the acetylase enzyme *α*TAT1 and the structural features of MTs, the patterns of acetylation formed on MTs are observed to be quite diverse. In this study, we present a multi-protofilament model with spatially heterogenous patterns of acetylation, and investigate how the local kinetic differences arising from heterogeneity affect the global kinetics of MT filaments. From the computational study we conclude that a filament with spatially uniform acetylation is least stable against disassembly, while ones with more clustered acetylation patterns may provide better resistance against disassembly. The increase in disassembly times for clustered pattern as compared to uniform pattern can be upto fifty percent for identical amounts of acetylation. Given that acetylated MTs affect several cellular functions as well as diseases such as cancer, our study indicates that spatial patterns of acetylation need to be focussed on, apart from the overall amount of acetylation.

**Author Summary:** Microtubules (MTs) form a crucial part of the cytoskeletal machinery which regulates several cellular processes. The basic building block of MTs are tubulin proteins. These proteins assemble in lateral and longitudinal directions to form a hollow cylindrical structure of a MT. There are chemical modifications on tubulin, known as Post Translational Modifications (PTMs), which affect the stability and dynamics of MT filaments. We computationally study how one such PTM, namely acetylation, affects the kinetics of disassembly of a MT filament. We propose a model which incorporates spatially heterogeneous patterns of acetylation on MT filament and study how they may regulate the disassembly times and velocities, a factor hitherto unexplored in studies. We conclude that there are significant differences of disassembly velocities and their fluctuations depending on the differnces in spatial patterns of acetylation.

## Introduction

Microtubules constitute an important class of bio-polymers that are essential for cell division, transport of vesicles, cell motility and for maintaining the structure and shape of the cell [1–5]. In order to fulfil certain cellular functions, during the corresponding stage of the cell cycle, MTs may have to polymerise and maintain stable structures. However, at other times, certain other cellular functions call for these filaments to depolymerise into free protein subunits, before entering into another period of stable assembly or bout of rapid dynamics. How MTs regulate the dynamics in these different states, and the switch between them, is an important question. It is known that certain chemical modifications on the polymer and various MT-binding proteins play crucial roles in regulating the stability and dynamics of the filaments. While there have been many models for investigating the dynamics of MTs alone, assembled *in vitro*, we know very little about how MT dynamics is influenced by the chemical modifications on the polymer. The focus of this work is to study the role of chemical modifications in deciding certain aspects of MT polymer dynamics.

The effect of these chemical modifications, known as Post-Translational Modifications (PTMs), on MT filaments depends on the structure and composition of the filaments. MT filaments exist as hollow cylinders of multiple protofilaments aligned laterally. Each protofilament consists of *α*-*β*-tubulin dimers (subunits). After tubulin subunits in a GTP-associated form polymerise to form MT filaments, the GTP molecule can undergo hydrolysis with a certain rate, to create GDP-associated tubulin subunits on the filament. In the absence of stabilizing external proteins, the higher depolymerisation rate of GDP-tubulin and the irreversibility of hydrolysis give rise to the phenomenon of “dynamic instability” in MTs, characterised by successive rapid shortening events (catastrophes) and slow growth events (rescues) [6]. The interaction of a variety of Microtubule Associated Proteins (MAPs) and PTMs on MT filaments may either alter the parameters associated with dynamic instability or altogether shift the filament into either a stable state which resists disassembly or an unstable state of enhanced disassembly [7].

There are a variety of enzymes which cause distinct modifications on tubulins such as acetylation, detyrosination, polyglutamylation, polyglycylation, polyamination etc. Tubulin subunits on a MT filament may exist in a variety of states with respect to the presence of each PTM. The extent to which each type of PTM is present on individual MTs correlates with the structure formed by these MTs, cell type and stages of the cell cycle [8, 9]. Mitotic spindle, centrioles and midbody show high levels of acetylation, detyrosination and polyglutamylation. Neurons also have high levels of these PTMs, with detyrosination being more abundant on the axons than growth cones. Axonemes, which form part of cilia and flagella, also have abundance of these PTMs, in addition to high levels of glycylation. However, PTMs are less abundant in astral MTs. Altering the amounts of PTMs in various cellular structures formed by MT filaments have been observed to affect their morphologies or functions to varying degrees [8, 10].

The exact mechanism by which each modification alters MT dynamics is not clearly understood. However, experimental evidences suggest that PTMs could influence MT dynamics by altering the interaction of other proteins with the filament [8, 11–13]. An alternate mechanism by which PTM can control MT dynamics is by changing inter-subunit interactions between adjacent protofilaments, as has been observed in the case of acetylation by recent *in vitro* experiments [14]. This is the first observation in which a PTM has been found to directly affect the lattice stability of MT filaments. In this work, we will focus primarily on acetylation that can alter the interaction between tubulin dimers and thereby change the kinetic behavior of growth/shrinkage.

Among the PTMs, acetylation has the peculiarity that it occurs at the lumen of the MT filament unlike most others which take place on the side chains of *α* or *β* tubulins. Primary position of tubulin acetylation is the Lys40 residue on *α* tubulin and the enzyme responsible is called *α*-tubulin acetyltransferase (*α*TAT1). *α*TAT1 has been shown to preferentially acetylate tubulins which are part of the MT filament rather than free tubulins in the solution [9]. If the lumen of the filament is presumed to be less accessible by proteins, aggravated by the decreased diffusion inside the cylinder [15], it can result in two scenarios. *α*TAT1 activity may be concentrated either at the open ends of the filament or at positions of defects in the lattice, through which they can presumably access the lumen or enter the tube. Despite this, *in vivo* experiments have observed diverse patterns of acetylation on cytoplasmic MT filaments in human fibroblast cells; the ensemble encompassed a large subpopulation of MTs with acetylation patches along their length and a small subpopulation with completely uniform distribution of acetylation on the MT lattice [16]. The peculiar position of acetylation and the resulting variety of acetylation patterns have become the subject of many ensuing *in vitro* experiments.

The idea that acetylation may be restricted near the position at which *α*TAT1 enters the lumen of the tube was corroborated by *in vitro* experiments which showed a gradient in acetylation from the tip towards the interior of MT filaments in axonemes [17]. However, later *in vitro* experiments have recorded a distribution of acetylation at random positions along the length of the filaments, without a preference for open ends [18]. Their findings favour a model in which acetylation is limited by low affinity of *α*TAT1 (high dissociation constant) for its substrate as well as a low catalytic rate. The low affinity enables the enzyme to diffuse to farther distances on the inner surface before catalysing acetylation, resulting in a more random pattern of acetylation along the filament length. This conclusion is in sharp contrast to the diffusion limited model of *α*TAT1 binding and acetylation observed by Coombes *et al* [19]. These *in vitro* experiments obtain acetylation patterns which are preferentially and abundantly distributed at the open ends of MTs. Their results support a model in which the mobility of the enzyme is low, which signifies high binding affinity and slow diffusion. These results are also supported by the experiments of Nathalie *et al* [15], which observed that *in vivo* MTs tend to have acetylation patches concentrated on the tips, whereas *in vitro* MTs showed random patches along the MTs. They attribute the presence of the random patches to defects along the filament and lateral entry of the enzyme through them.

These set of experiments suggest to us that *several acetylation patterns can be formed* on MT lattices depending on filament geometry as well as the binding and catalytic rates of the acetyltransferase. Given that this variability also exists in the case of *in vivo* MT filaments [16], it would be interesting to see *whether these patterns have a direct impact in regulating the dynamics of MTs*.

To understand the effect of acetylation on the stability of MT lattice, it is essential to know how this modification affects the mechanical and kinetic parameters of MTs. There have been some earlier experimental studies which mapped the overall changes in MT population caused by altering the amount of acetylation. Experiments on touch receptive neurons (TRN) of C. elegans [20, 21] observed that suppression of acetylation on MTs results in shorter filaments with protofilament number varying between 10-16, whereas, the wild-type cells mostly contain long MT filaments with 15 protofilaments. Recent experiments using cryo-electron microscopy [22] probed potential differences in MT structure caused by acetylation. They observe that the state of tubulin acetylation does not cause any discernible variation in the helical structure of MTs, retaining the same lateral and longitudinal spacing in acetylated and deacteylated MTs. Neither do they find significant differences in the structure of the tubulin dimer at a fairly high resolution of 8 – 9*Å*.

However, recent controlled experiments by Portran *et al* [14] had, for the first time, revealed how the effect of acetylation is manifested in the physical properties of individual filaments. Their results show that the effect of acetylation on MT lattice is to (i) *reduce the strength of lateral interaction* between tubulin subunits on adjacent protofilaments. This is observed in direct FRET and negative-stain electron microscopy measurements as well as discerned from the reduced nucleation and self assembly rates of MT filaments formed from acetylated tubulin dimers compared to the deacetylated ones. In addition, from single MT growth trajectories, they find that (ii) *acetylation increases the rate of shrinking in MTs threefold* while maintaining the rate of elongation at a constant value. Also, (iii) *there is no significant effect of acetylation on catastrophe rates*.

In this work we would base our model on the experimental results by Portran *et al* [14]. The effects of acetylation on microtubule filament as explained in Portran *et al* [14] are drawn from experiments conducted with either fully acetylated or fully deacetylated MTs. Instead, if specific patterns of acetylation on MT lattice are considered, those effects would mean that acetylated tubulins can form unstable regions or domains of different shapes and sizes on the lattice.

Variability in acetylation patterns has been observed in MTs assembled under different conditions in both *in vivo* and *in vitro* experiments. In this study we investigate to what extent can the phenomenological parameters such as disassembly times and velocities be regulated by distinct patterns of acetylation. Our results show that the profiles of phenomenological parameters associated with filament disassembly can shed light on the underlying configuration of acetylation. In the following section, a multi-protofilament model is explained, which is capable of accommodating the various patterns or distributions of acetylation on the MT lattice, as well as their effect on inter-protofilament interactions.

## Model and Methods

Theoretical studies model MT filaments in terms of abstract objects which interact via various physical laws, in order to explain their complex dynamics observed in experiments. The models differ among themeselves in terms of the extent of coarse-graining and the type of parameters used in them [23]. One set of extensive coarse-grained theoretical models that study MT dynamics, use the four phenomenological parameters associated with “dynamic instability”. These minimal set of parameters are the mean velocities of growth and shrinkage and the frequencies of catastrophes and rescues [24]. Another set of models explicitly include microscopic kinetic rates associated with processes such as polymerisation, depolymerisation and hydrolysis [25–29]. This type of models are coarse-grained to some extent since the knietic rates (whose values are measured from experiments) are input parameters rather than emerging from the mechanical and chemical properties of tubulin proteins. However, the inclusion of kinetic rates of various reactions associated with individual tubulin subunits does allow for probing the physical basis of the emergence of many macroscopic dynamic features of the MT filaments. For example, in an earlier work, using a linear polymer model, we observe that the differential polymerisation rate at the open MT tip containing a GTP-tubulin versus a GDP-tubulin can regulate the multi-step mechanism behind age-dependent catastrophes [30]. In another category of models, the multi-protofilament structure of MT filaments, which may include bond energies between subunits and elastic properties of protofilaments, is accounted for in detail [31–39]. The choice of employment of the level of coarse-graining in theoretical models depends on the aspect of MT dynamics that they seek to answer.

It is known that acetylation alters the lateral interaction between tubulin subunits on adjacent protofilaments and thereby the kinetics of these subunits [14]. Hence, given that our aim in this paper is to study how the spread of acetylation influences the dynamics of MTs, it may be suitable for us to employ a multi-protofilament model that has microscopic kinetic rates of polymerisation and depolymerisation which depend on the interaction between subunits on lateral protofilaments. To acheive this we start with a model similar to that presented in Stukalin & Kolomeisky [40, 41] and VanBuren *et al* [31], and extend it further to include the effect of chemical modifications. The detailed features of the model and the accommodation of acetylation are explained in the following subsections.

### A multi-protofilament lattice model to study MT stability

In the model we employ, MT filament is represented by an *N*-protofilament lattice (with *N* = 13), as shown in the schematic (Fig 1(a)). Each *α*-*β*-tubulin dimer is represented by a single subunit with lateral and longitudinal neighbours. Periodic boundary conditions were used such that subunits on protofilament numbers 1 and *N* are lateral neighbours. One end (on the left of Fig 1(a)) of the filament is open where polymerisation (depolymerisation) can occur via the association (dissociation) of individual subunits to (from) any protofilament. The other end (on the right of Fig 1(a)) is assumed as a stable nucleated seed which acts as a reflecting boundary.

**Fig 1.**
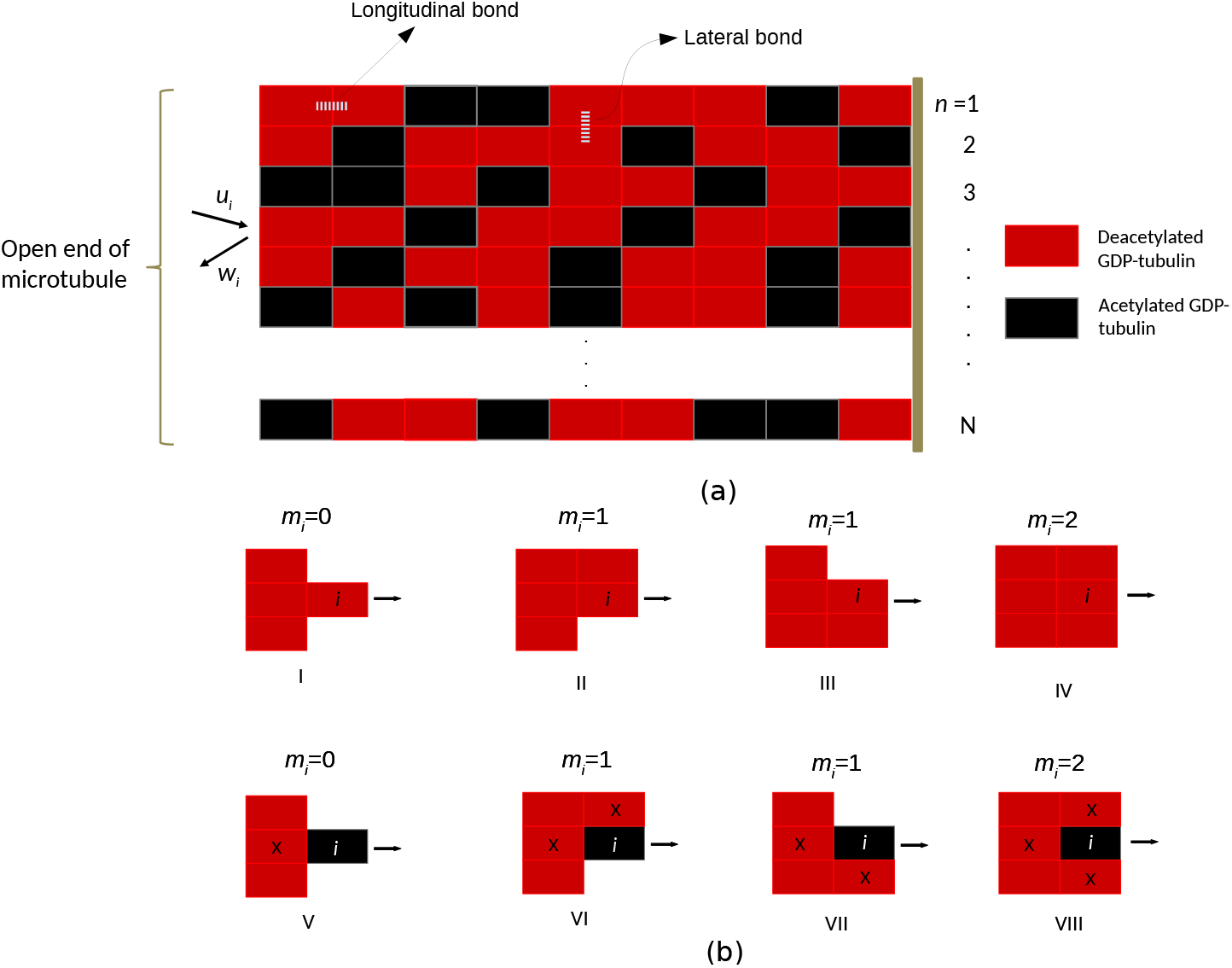
(a) The 13-protofilament MT cylindrical structure is approximated with a multi-protofilament lattice with 13 linear filaments, with periodic boundary such that filaments 1 and 13 are neighbours. Polymerisation and depolymerisation at the *i^th^* subunit take place with rates 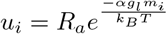 and 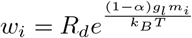, respectively. The red blocks represent deacetylated GDP-tubulins and the black blocks represent acetylated GDP-tubulins. (b) The rates of polymerisation and depolymerisation depend on the fraction of lateral bonds the subunit forms with the neighbours (*m_i_*). In the simplistic case where the shifts between protofilaments are integer multiples of one tubulin subunit length, *m_i_* can take values 0, 1 and 2. In the figure, I-IV represent the depolymerisation of *i^th^* subunit in deacetylated state and V-VIII represent the depolymerisation of *i^th^* subunit in acetylated state. Corresponding values of *m_i_* that determine the rate of depolymerisation are written alongside the schematic. Note that 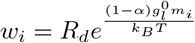 in I-IV. Whereas, in V-VIII, 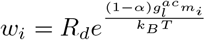. In addition, note that, when any of the nearest neighbours of *i* (marked → ‘x’) is acetylated, this also corresponds to 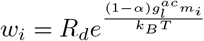.

In cell, acetylation has been observed to predominantly exist on long-lived MTs stablised by other proteins, and the time-scales of acetylation observed in bulk measurements (≈ 0.03min^−1^) [18] are much longer than those of GTP-hydrolysis (≈ 20min^−1^) [2]. Hence, we speculate that in cells the presence of acetylation patterns may be more prevalent on a lattice consisting of only GDP-tubulins. Therefore, their effect on a GDP-tubulin-lattice may be more biologically significant. The *in vitro* experiments of Portran *et al* [14] also show the effect of acetylation on shrinkage rate of MTs undergoing catastrophes, during which they mostly consist of GDP-tubulins. Hence, in this study, the filament lattice consists of GDP-tubulins, on which acetylation can be thought to have introduced local spatial disorders. Experimental evidences show that at these points of disorder, lateral interaction energy between subunits of neighbouring protofilaments is reduced [14]. Hence, in the model, the lateral interaction energy (*g_l_*) between subunits can assume two values; 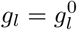 in the absence of acetylation, and the points of disorders introduced by acetylation are characterised by an altered value of lateral interaction energy represented by 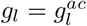.

Consider a filament of length *L*. Let *a_i,n_* represent the state of acetylation of the subunit on protofilament number *n* at position *i* where *i* = 1 (closed end) to *L* (open end), and *n* = 1 to 13. Then,

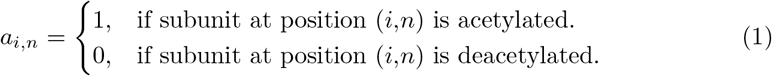

The value of lateral interaction energy (*g_l_*) associated with the subunit at (*i,n*) depends on the status of acetylation of subunits at (*i,n*), (*i* – 1, *n*), (*i, n* – 1) and (*i, n* + 1). Hence, we focus on the set *I* = {*a_i,n_*, *a*_*i*–1, *n*_, *a*_*i, n*–1_, *a*_*i,n*+1_} of indicator variables. In order to determine the rates of polymerisation and depolymerisation at position (*i,n*), the lateral interaction energy shared with the neighbours is determined as,

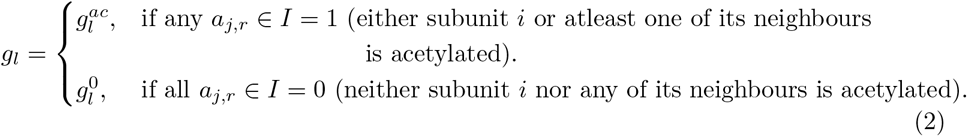

The rates of polymerisation and depolymerisation of a subunit at position (*i,n*) (on the left of Fig 1(a)) are dependent on the local configuration around the subunit, in addition to the value of *g_l_* associated with it. The rates of polymerisation and depolymerisation are *R_a_* and *R_d_*, respectively, when the subunit has no lateral neighbour. For all the other configurations, rate of polymerisation is given by 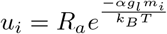. *R_a_* is linearly proportional to the concentration of free tubulin. However, since our focus in this paper is on the disassembly profiles of the filament, we simulate the filament in the absence of free tubulin in the solution. Hence, *u_i_* =0 throughout the study, while the rate of depolymerisation is given by,

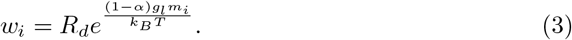

Here *m_i_* is the number of lateral neighbours shared by the subunit at position *i* which can take values 0,1 or 2 (see Fig 1(b)). The constants *α* and 1 – *α* represent the relative fractional contribution of the lateral interaction energy to the rates of polymerisation and depolymerisation, respectively. We use kinetic Monte Carlo simulations to model the kinetics of the system [42]. The parameters and their values used in the simulations are listed in Table 1 (A detailed description of how their values are decided from the information available in earlier computational and experimental studies is provided in Supporting Information (SI)).

**Table 1.**
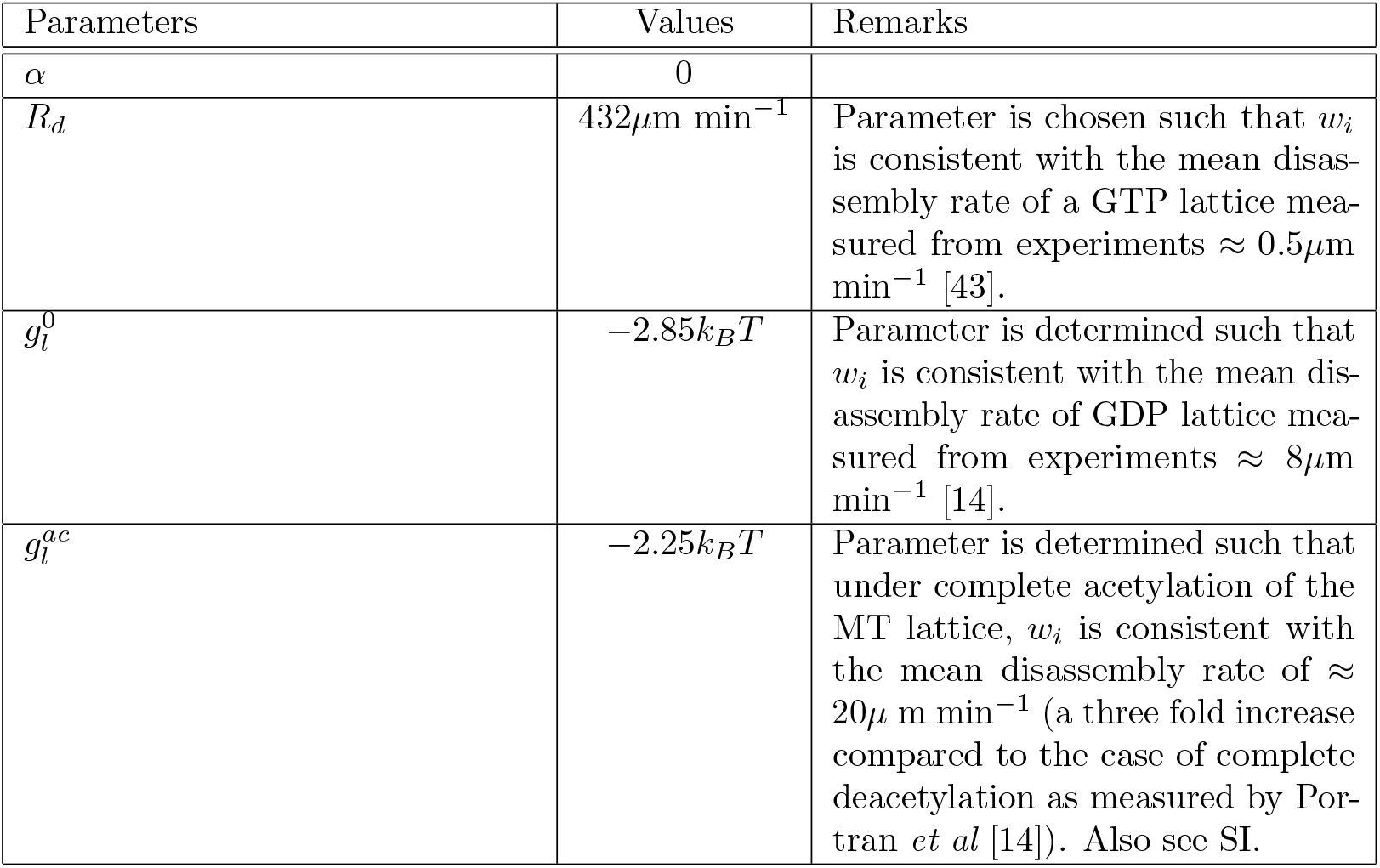
Parameters used in the simulations

| Parameters | Values | Remarks |
| --- | --- | --- |
| $\alpha$ | 0 | |
| $R_d$ | $432\mu\text{m min}^{-1}$ | Parameter is chosen such that $w_i$ is consistent with the mean disassembly rate of a GTP lattice measured from experiments $\approx 0.5\mu\text{m min}^{-1}$ [43]. |
| $g_l^0$ | $-2.85k_B T$ | Parameter is determined such that $w_i$ is consistent with the mean disassembly rate of GDP lattice measured from experiments $\approx 8\mu\text{m min}^{-1}$ [14]. |
| $g_l^{ac}$ | $-2.25k_B T$ | Parameter is determined such that under complete acetylation of the MT lattice, $w_i$ is consistent with the mean disassembly rate of $\approx 20\mu\text{m min}^{-1}$ (a three fold increase compared to the case of complete deacetylation as measured by Portran *et al* [14]). Also see SI. |

In this study, we consider completely hydrolysed MT filaments consisting of only GDP-tubulin subunits. Information available from experiments shows that various acetylation patterns can be formed on stable MT lattices which can potentially arise from differences in the mechanism of *α*TAT1 entry, diffusion and catalysis. In this work, we incorporate distinct preformed acetylation patterns onto the GDP-tubulin lattice, in accordance with these earlier experimental observations. At various points in the cell cycle, a stable filament is required to undergo rapid disassembly before the next assembly event begins [16]. There are also various dilution assay experiments which study the disassembly dynamics of filaments. A combination of these factors serve as motivation for us to study the disassembly of MT filaments with various patterns of acetylation, when the free tubulin concentration in the solution and, therefore, the rate of polymerisation are zero.

### Patterns of acetylation on the MT lattice

Acetylation patterns formed on a stable MT depend on the mechanism of the TAT entry, luminal diffusion of the enzyme on the filament surface, and its catalysis rate. Based on *in vitro* experimental results which broadly take into account these factors, we have marked the MT lattice with various patterns of acetylation. *In vitro* experiments have observed that the effect of acetylation is to reduce lateral interaction between subunits. As a result, the accumulation of various acetylation patterns is equivalent to accumulation of corresponding unstable regions on the filament lattice. Hence, it will be of interest to learn, in the event of disassembly, what role do these patterns have in regulating the stability of MTs. Fig 2(a)-(c) show three distinct types of patterns of acetylation on the filament. On the left, patterns of acetylation on the lattice are shown with “red” representing deacetylated tubulin subunits and “black” representing acetylated tubulin subunits. On the right, the patterns are quantified with the *l*-axis corresponding to the length (in *μ*m) measured from the tip of the filament towards the interior of the filament (for an equivalent length as the corresponding figure on the left). The quantity *f*(*l*) in the figure represents the fraction of protofilaments in acetylated state (out of the 13 protofilaments) for the corresponding length *l*.

**Fig 2.**
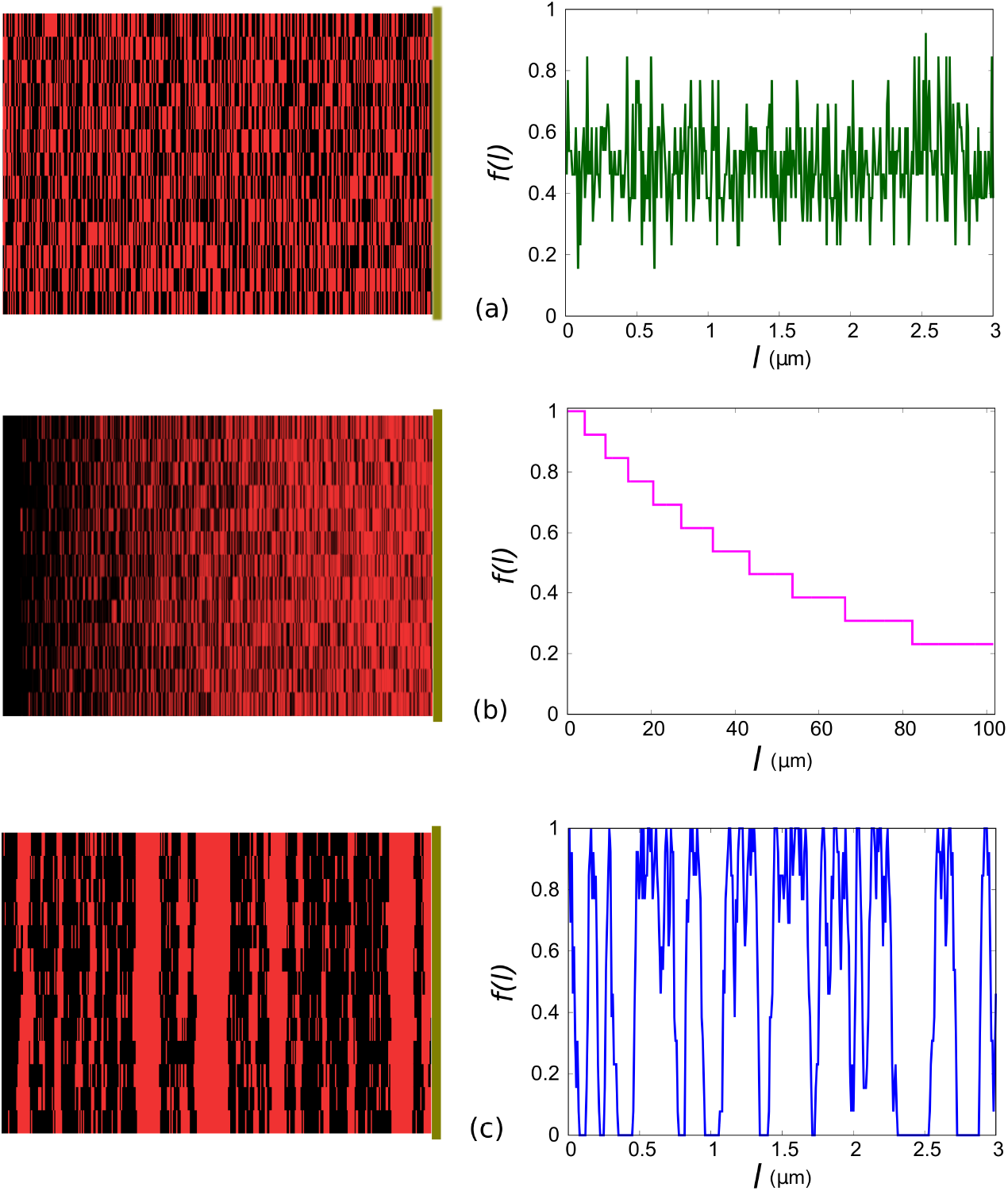
Left: Uniform (a), exponentially decaying (b) and clustered (c) acetylation patterns are represented on the filament lattice with “red” representing deacetylated tubulin subunits and “black” representing acetylated tubulin subunits. Right: Fraction of protofilaments acetylated per layer (*f*(*l*)) as a function of the length (*l*) of the MT filament, correspoding to patterns in left panel of (a), (b) and (c). The fraction of acetylation *ρ_ac_* = 0.5, is the same for all three curves. In (c), the fraction of defect layers *ρ_ld_* = 0.1. See text for definition of *ρ_ld_*.

An important parameter to consider is the total fraction of acetylation *ρ_ac_*, defined as the number of acetylated subunits on the filament divided by the total number of subunits on the filament. In Fig 2(a)-(c), *ρ_ac_* = 0.5. Note that when the pattern of acetylation is clusterd (Fig 2(c)), there is an additional parameter which needs to be considered which is represented by *ρ_ld_*. This parameter represents the fraction of positions, along the axis of the filament, on which a defect is assumed to be present. Defects are MT lattice openings through which the TAT enzyme may enter the luminal space. Note that *ld* stands for defect layer. For the clustered pattern, likelihood of acetylation is maximum at defect layers. In other words, these are the layers at which maximum number of subunits are acetylated. Descrition of three distinct patterns of acetylation considered in this study are provided in the following paragraphs.

#### Uniform acetylation pattern

Fig 2(a) corresponds to a uniform pattern of acetylation, in which acetylation can occur on any random subunit on the MT filament with equal probability without a preference for any position. This is in accordance with a situation where the enzyme can enter the lumen of the filament either through the open tip or through any defect (lattice opening) along the filament, or both. After entering, high diffusion constant and small catalysis rate of the enzyme cause acetylation pattern to be completely uniform.

#### Exponential acetylation pattern

Fig 2(b) corresponds to the exponential pattern of acetylation. This can arise in a scenario where the enzyme enters the lumen of the filament only through the filament tip, and due to slow diffusion compared to the catalysis rate, acetylation proceeds with a gradient from the tip towards the interior. As a result, acetylation begins at the tip of the filament and decreases exponentially towards the interior of the filament. In each of the subunit layer along the length of the filament, however, acetylation is random and can take place on subunits on any of the 13 protofilaments with equal probability. The spatial pattern follows the form *exp*(−*bl*), where *l* is the length of the filament measured from the open tip. At the open tip (*l* = 0), subunits on all 13 protofilaments are acetylated. Additionally, the curve has discrete steps since the protofilament numbers and filament length, when expressed in terms of subunits, are integers. The parameter *b* is determined for every value of *ρ_ac_*, and is dependent on the length of the filament.

#### Clustered acetylation pattern

Fig 2(c) corresponds to a clustered acetylation pattern. In this case, the enzyme may enter through both the tip as well as defects on the lattice. Defects can be present at any random layer along the length of the filament. A lateral layer containing atleast one defect is called a defect layer. Acetylation can commence from any of the defect layers. Since, acetylation also commences from the open tip, it is also one of the defect layers. In Figure 2(c) blue curve, the fraction of defect layers, *ρ_ld_* is 0.1. i.e., ten percent of the layers along the filament are such that acetylation commences from those. The clustered pattern can potentially arise from two scenarios. In the first scenario, luminal surface diffusion of the enzyme is slow compared to the catalysis rate. As a result, acetylation takes place near the position of entry before the enzyme diffuses away on the luminal surface. In the second scenario, the enzyme has a high rate of exit through the defect through which it entered. In this case too, the most probable position for acetylation is near the position of entry. In our model, the clustered acetylation pattern is generated according to the following procedure. If the fraction of total acetylation is equal to or greater than the fraction of defect layers (*ρ_ac_* ≥ *ρ_ld_*), the defect layers undergo complete acetylation (subunits on all 13 protofilaments are acetylated). In the case of *ρ_ac_* > *ρ_ld_*, for further acetylation of the remaining (*ρ_ac_* – *ρ_ld_*) × *N* × *L* subunits (where *N* is the protofilament number and *L* is the filament length in subunits), a subunit in the whole filament is chosen at random and it is acetylated if any one of its four nearest neighbours is acetylated. This procedure continues until the total number of acetylated subunits reaches the number = *ρ_ac_* × *N* × *L*. Note that since the state of acetylation of a subunit depends on those of its neighbours, acetylated clusters arise around the defect layers. In order to form acetylation patterns in the case of *ρ_ac_* < *ρ_ld_*, a minimum of 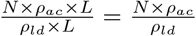 subunits are acetylated at each defect layer. Note that the positions of these subunits along that particular layer are chosen randomly. For further acetylation, a subunit is chosen randomly and it is acetylated if it belongs to a defect layer. As before, this procedure continues until the total number of acetylated subunits reaches the number stipulated by *ρ_ac_*.

## Results

We use kinetic Monte-Carlo [42] simulations on the multi-protofilament model and generate length versus time data during disassembly. Using these data, distributions of disassembly times and velocities are measured for an ensemble of 3 × 10^4^ filaments. Stability of the filaments with various patterns of acetylation are explored using statistical quantities measured from the corresponding distributions, as explained in the following subsections.

### Total disassembly times are regulated by acetylation patterns

In this section, we investigate whether the time it takes for an ensemble of filaments to convert from the polymeric form to the dimeric form be regulated by the inclusion of different patterns of acetylation. In Fig 3(a)-(b), length of the filament (measured as the length of the longest protofilament at every step) is plotted as a function of time as the disassembly progresses. The pattern of acetylation is uniform in Fig 3(a) and (b). In Fig 3(a), the two curves show distinct paths of disassembly followed by the filament starting from identical patterns of acetylation. The differences arise due to the stochasticity inherent in the kinetics of the process due to thermal fluctuations. In kinetic Monte-Carlo, the times of every next event in the process is drawn from an exponential distribution and the events themselves are chosen with a probability proportional to their kinetic rates. In Fig 3(b), the purple curve is retained from (a). However, the black curve corresponds to a filament where the positions of acetylated subunits have been altered, although the overall pattern is still uniform. Note that, in this figure, the differences between the paths followed by the two curves arise due to the spatial stochasticity associated with the distribution of disorders of acetylation across the filament lattice. In reality, stochasticity in the length versus time data arises due to both these factors – the spatial variation of acetylation as well as thermal fluctuations. All the measurements in this study, hence, invoke averages over these two sources of randomness.

**Fig 3.**
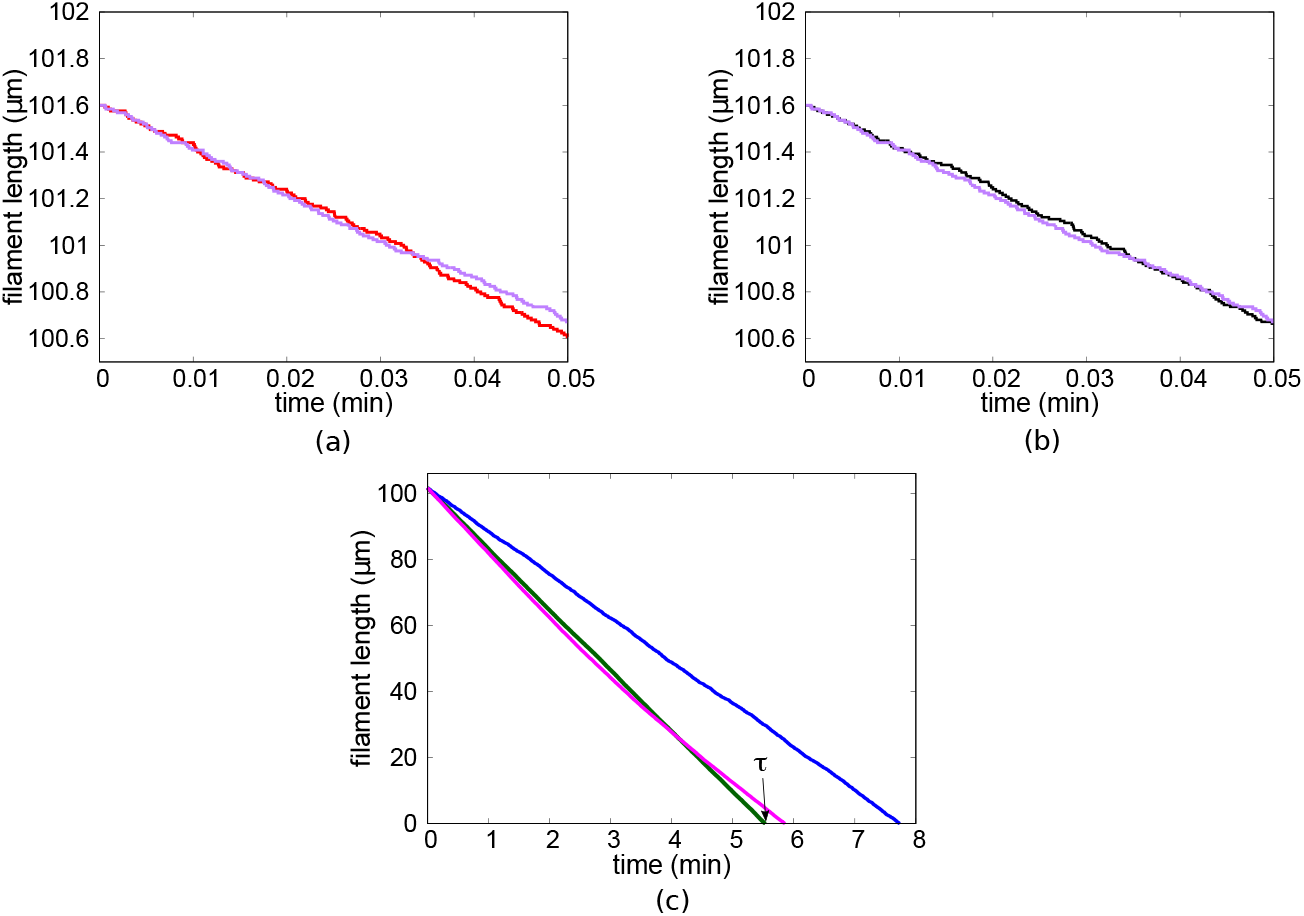
(a)-(b) show length of the filament (measured as the length of the longest protofilament at every step) is plotted as a function of time, as the filament, with uniform acetylation pattern, undergoes disassembly in the presence of *ρ_ac_* = 0.5. (a) The two curves (red and purple) correspond to two realisations, where the difference between the two curves arises due to the stochasticity associated with the kinetics of the process, despite the measurements being done on filaments with identical disorder patterns introduced by acetylation. (b) Length versus time data corresponding to two realisations (black and purple) show stochasticity arising due to the difference in disorder patterns (positions of acetylated subunits) across the lattice. (c) Length versus time data for uniform (green), exponential (magenta) and clustered (with *ρ_ld_* = 0.1 (blue)) acetylation patterns. Total disassembly time is represented by *τ* (as shown for the uniform case by arrow near the green curve), measured for disassembly from an initial length of ≈ 102*μ*m to zero length.

Fig 3(c) presents the comparison of the length versus time data under uniform (green), exponential (magenta) and clustered (with *ρ_ld_* = 0.1 (blue)) acetylation patterns. The variation in filament stability can be assessed from the total time of disassembly for the filament to shrink from a finite length (≈ 102*μ*m in this study) to zero length. The longer it takes to disassemble the more stable the filament is. Total disassembly times are represented by *τ* (indicated in Fig 3(c) for the uniform pattern, green). Our first important result is that *τ* corresponding to exponential and uniform patterns are shorter compared to that of the clustered pattern with *ρ_ld_* = 0.1.

Note that due to the stochasticities associated with the disassembly of the filament, *τ* is a random variable, and for different realisations, it takes different values. Hence, in order to compare the disassembly behaviour under various patterns, we may find the *mean* and other moments of *τ*. A comparison of disassembly times for a range of *ρ_ac_* are plotted in Fig 4(a). The curves show the *mean* total disassembly time 〈*τ*〉 as a function of *ρ_ac_* for all types of patterns discussed earlier. The *mean* total disassembly times for clustered acetylation pattern (for *ρ_ld_* = 0.1 (blue)) are in general larger than those for both uniform (green) and exponential (magenta) acetylation patterns, for all *ρ_ac_* near and greater than the corresponding *ρ_ld_* value (for example, near and greater than *ρ_ac_* = 0.1 in the blue curve). In Fig 4(b), a comparison of mean disassembly times corresponding to different clustered patterns of acetylation, with *ρ_ld_* = 0.1 (blue), *ρ_ld_* = 0.2 (black) and *ρ_ld_* = 0.3 (brown), are plotted. It may be expected, as *ρ_ld_* approaches 1, the curves for clustered pattern approach the one corresponding to the uniform pattern.

**Fig 4.**
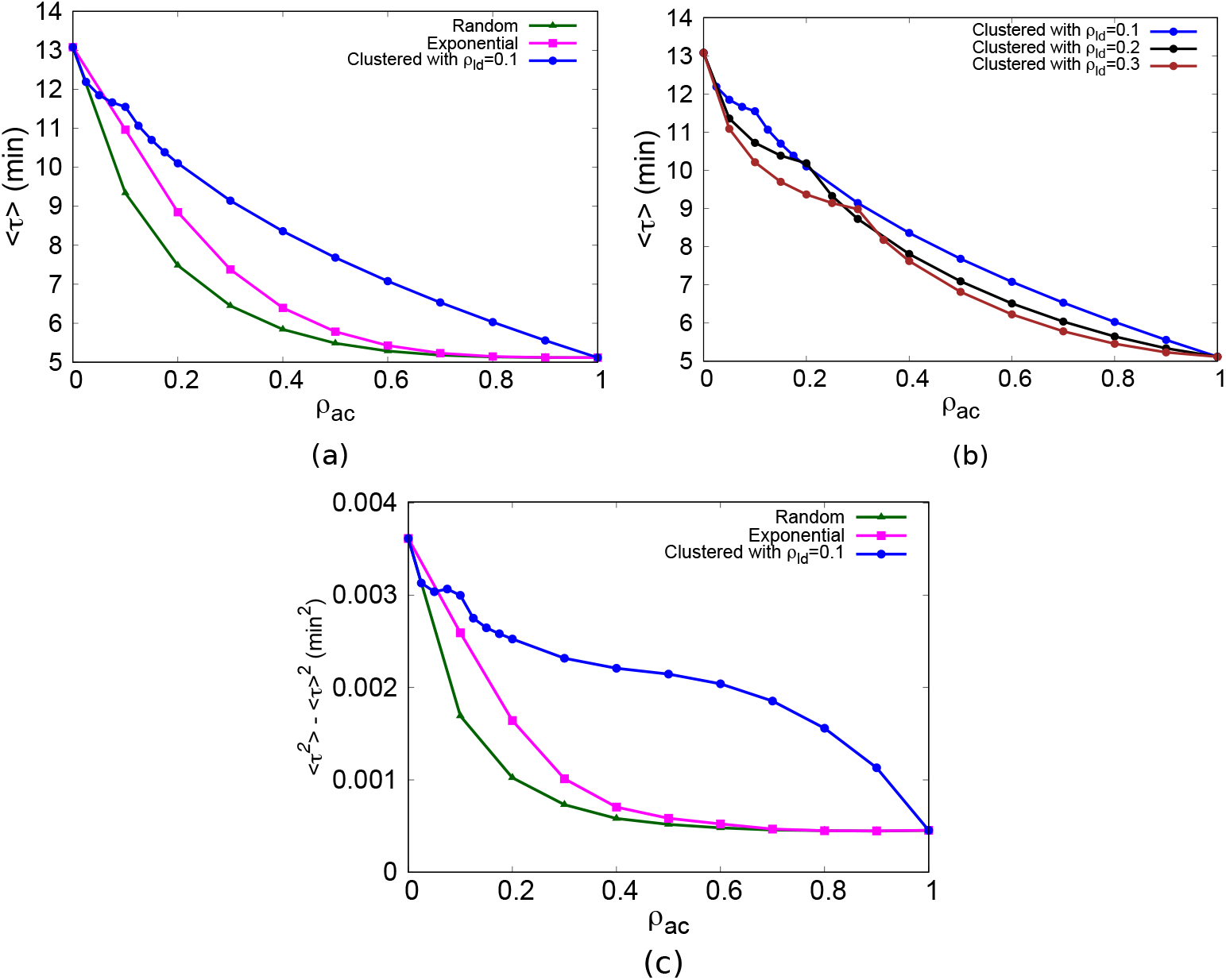
(a) *Mean* of *τ* (〈*τ*〉) is plotted as a function of total fraction of acetylation *ρ_ac_* for uniform (green), exponential (magenta) and clustered (with *ρ_ld_* = 0.1 (blue)) patterns. In (b), the curves show 〈*τ*〉 as a function of *ρ_ac_*, for different clustered patterns with *ρ_ld_* = 0.1 (blue), *ρ_ld_* = 0.2 (black) and *ρ_ld_* = 0.3 (brown). In (c), *variance* of *τ* (measured as 〈*τ*^2^〉 - 〈*τ*〉^2^) of filaments is plotted as a function of *ρ_ac_* for uniform (green), exponential (magenta) and clustered (with *ρ_ld_* = 0.1 (blue)) patterns.

Since *τ* is a random variable, it is associated with fluctuations, which may be quantified through the variance of the distribution of *τ*. In Fig 4(c), the *variance* of *τ* (measured as 〈*τ*^2^〉 - 〈*τ*〉^2^) is plotted as a function of *ρ_ac_*. In the case of uniform (green) and exponential (magenta) patterns, the *variance* monotonically decreases with increase in *ρ_ac_* until it reaches a saturation value for higher values of *ρ_ac_*. This can be contrasted with the *variance* curve of the clustered pattern (blue), where the *variance* initially decreases, followed by a saturation near *ρ_ld_* = *ρ_ac_*. Thereafter, the *variance* follows a slow decrease for higher values of *ρ_ac_*. The nature of each of these curves is dictated by the internal clustering of acetylated subunits on the MT lattice for the corresponding pattern.

### Disassembly velocities characterise the underlying acetylation pattern

We calculate disassembly velocities (*v*) for individual filaments, over an interval of every 1/60 min, from the length versus time data corresponding to various realisations, for a given value of *ρ_ac_* and *ρ_ld_*. In Fig 5(a), *v* obtained from one realisation is plotted as a function of time, for uniform pattern of acetylation (green) and clustered pattern of acetylation with *ρ_ld_* = 0.1 (blue), with *ρ_ac_* = 0.5 in both cases. Since, *v* is a stochastic variable, each realisation gives rise to a distinct trajectory of *v* as a function of time. The probability distribution of *v* calculated from velocity vs time curves, from an ensemble of 3 × 10^4^ filaments (or realisations), is plotted in Fig 5(b). Here, each *v* is measured at a fixed time ≈ 1.5 min. The vertical lines in the figure correspond to the respective *mean* values (〈*v*〉) of the distributions. While the uniform pattern has a higher *mean* velocity compared to the clustered pattern, the latter has a much wider distribution compared to the former.

**Fig 5.**
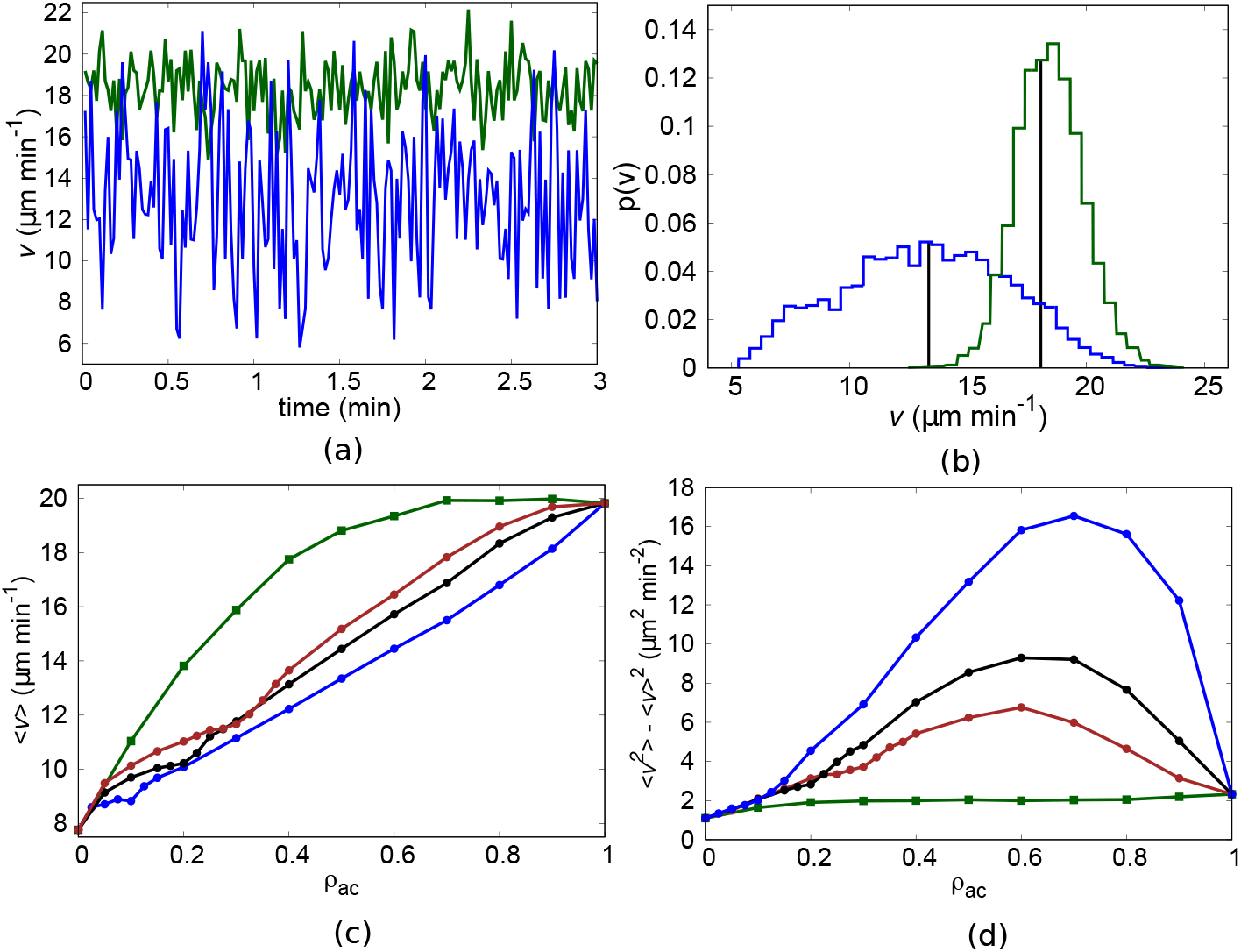
(a) Velocity of disassembly (*v*) of filaments (measured for every interval of (1/60) min from the length versus time data) for uniform acetylation pattern (green) and clustered acetylation pattern with *ρ_ld_* = 0.1 (blue) is plotted as a function of time. Total fraction of acetylation *ρ_ac_* = 0.5. In (b), the corresponding distribution of velocities (measured at time ≈ 1.5 min) is plotted, with the vertical lines marking the mean value (〈*v*〉). (c) 〈*v*〉 is plotted for a range of fractions *ρ_ac_* for uniform pattern (green) and clustered patterns with *ρ_ld_* = 0.1 (blue), 0.2 (black) and 0.3 (brown). (d) Variance of *v*, calculated as 〈*v*^2^〉 - 〈*v*〉^2^ from the distribution, is plotted as a function of *ρ_ac_* for uniform and clustered patterns.

In Fig 5(c), 〈*v*〉 values calculated from the probability distributions are plotted as a function of the corresponding *ρ_ac_* values. The uniform acetylation pattern (green) has a distinctly higher 〈*v*〉 compared to the clustered acetylation patterns (blue, black, brown) for all the intermediate *ρ_ac_*. Among the clustered patterns, the one corresponding to the least number of defect layers (*ρ_ld_* = 0.1, blue) constitutes the most stable lattice against disassembly.

The difference in disorder created on the lattice due to acetylation patterns not only affects the mean of *v*, but strongly alters the fluctuations associated with *v* as well. This is expressed as the *variance* of *v* (calculated as 〈*v*^2^〉 - 〈*v*〉^2^) plotted in Fig 5(d) as a function of *ρ_ac_*, corresponding to uniform and clustered acetylation patterns. For all *ρ_ac_*, *variance* is the least for uniform acetylation pattern (green). Among the clustered acetylation patterns, *ρ_ld_* = 0.1 (with least number of defect layers) has a distinctly higher *variance* compared to others.

Next, we verify the disassembly profile of the filament with exponential acetylation pattern. Fig 6 shows the *mean* velocity (black) expected from an exponential acetylation pattern (magenta) plotted as a function of distance from the tip, for *ρ_ac_* = 0.5. The exponential pattern of acetylation consists of fully acetylated layers at the open end of the filament, followed by a decay in the extent of acetylation according to the form *exp*(−*bl*), where *l* is the length of the filament measured from the open tip. Hence, if *v*_0_ is the *mean* disassembly velocity of the filament in the absence of any acetylation (≈ 7.76*μ*m min^−1^ in this study) and *v*′ + *v*_0_ is the maximum *mean* disassembly velocity in the presence of complete acetylation at *l* = 0*μm* (≈ 19.81*μ*m min^−1^ in this study), then the mean velocity is expected to follow the form *v*′ *exp*(−*bl*) + *v*_0_ as l increases (Fig 6 (black)). However, the 〈*v*〉 of the filament calculated from simulations (red) are much higher than the expected velocities along the length of the filament. This decreased stability of the filament can be attributed to acetylated subunits being distributed randomly along each layer. As a result, even when only a few subunits on each layer are acetylated, the decrease in their lateral interaction energies ensures that their neighbours also (and by extension the layer itself) have less resistance against disassembly. This cooperative behaviour is discussed in detail below.

**Fig 6.**
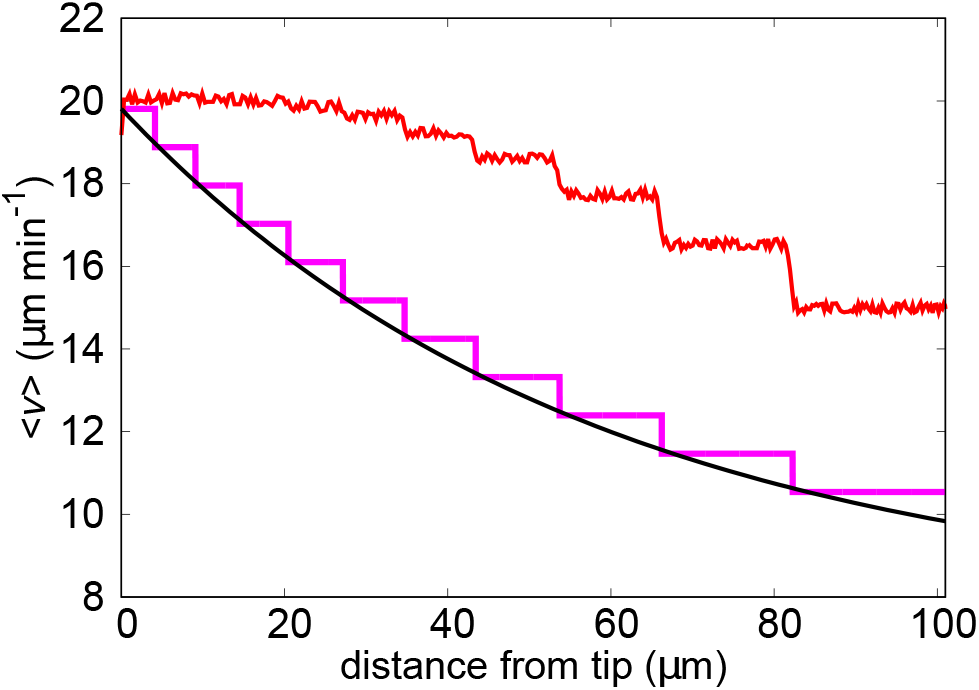
Mean velocities expected (black) from the exponential acetylation pattern are compared with mean velocities calculated (measured for an interval of (1/60) min from the length versus time data) from simulations (red) performed with the exponential acetylation pattern, both plotted as a function of the distance from the open tip of the filament, for *ρ_ac_* = 0.5. The larger values of mean disassembly velocities obtained from simulations (red) arise as a result of decresed stability caused by random distribution of acetylated subunits along each lateral layer of protofilaments. A scaled exponential pattern of acetylation (magenta) is plotted as a marker.

### Cooperativity on a lateral layer affects the observed kinetics

Acetylation decreases the lateral interaction energy between subunits of neighbouring protofilaments. Hence, the effect of various patterns of acetylation on filament disassembly depends on the number and proximity of acetylated subunits on each lateral layer. Number of acetylated subunits per layer (*k*) can vary from 0 to 13 (number of protofilaments), and is plotted along the x-axis of Fig 7. Y-axis in Fig 7 corresponds to the fraction of lateral layers, in a MT filament, with *k* acetylated subunits – sampling is done over an ensemble of multiple filaments. Here, *ρ_ac_* = 0.5 for both uniform acetylation pattern (green) and clustered acetylation pattern (blue).

**Fig 7.**
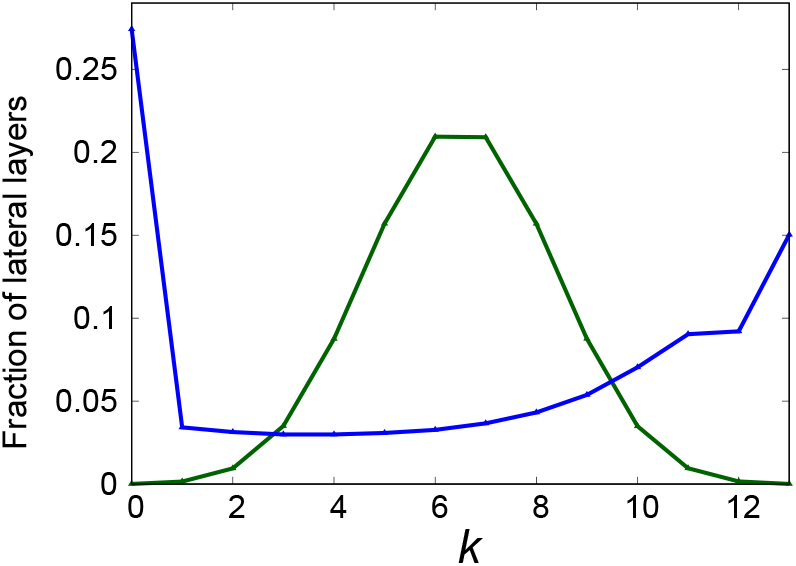
Fraction of lateral layers (rings) per MT filament, which have *k* ∈ [0,13] acetylated subunits is plotted in the y-axis as a function of *k* in the x-axis. Here, *ρ_ac_* = 0.5 and the curves correspond to uniform pattern (green) and clustered pattern with *ρ_ld_* = 0.1 (blue). For uniform pattern, fraction is unimodal and the peak appears at *k* = 6 ≈ *N ρ_ac_*, where *N* = 13. For the clustered pattern, fraction is bimodal; the peak at *k* = 13 arises due to the assumption that each defect layer is completely acetylated when *ρ_ac_* ≧ *ρ_ld_* and the one at *k* = 0 signifies the large number of completely deacetylated lattice layers.

The stability of every lateral layer depends on the fraction and relative positions of acetylated subunits in them. Since, the effect of acetylation is to reduce the lateral interaction energy between subunits on neighbouring protofilaments, it follows that in a lateral layer, a deacetylated subunit has a reduced lateral interaction energy 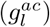 when atleast one of its neighbours is acetylated. When a minimum of approximately half the layer is acetylated (*k* = 6), this influence from acetylated neighbours gives rise to a cooperative decrease in lateral energy and increase in local disassembly velocity of the entire lateral layer. On the other hand, lateral layers which contain less than *k* = 6 acetylated subunits have higher chances of containing three or more consecutive deacetylated subunits. This is marked by a decrease in neighbour-induced cooperative disassembly at those layers. Hence, the local disassembly velocity at these layers is smaller compared to the former case.

In the case of uniform pattern, the randomness in the distribution of acetylated subunits over the MT lattice causes many lateral layers (rings) to contain approximately half the number of protofilaments in acetylated state in Fig 7 (green) (*k* ≈ *N ρ_ac_* = 13 * 0.5 ≈ 6). Hence, the fraction in Fig 7(green) is unimodal and the peak is at *k* = 6. Here, the total fraction of lateral layers with *k* ≥ 6 is 0.7, and that with *k* < 6 is 0.3. Hence, according to the arguments presented above, there are substantially more number of layers which have larger values of local disassembly velocities. This results in the larger *mean* disassembly velocities of filaments with uniform pattern as seen in Fig 5((c), green).

This can be contrasted with clustered patterns (for *ρ_ac_* ≥ *ρ_ld_*), where, a large number of layers are completely acetylated (*k* = 13), giving rise to a considerable number of completely deacetylated layers (*k* = 0) and a smaller number of layers with intermediate values of *k*. This causes the occurence of bimodality of fraction in Fig 7(blue, with *ρ_ld_* = 0.1). The peaks at *k* = 0 and *k* = 13, are wide apart from each other, and correspond to completely deacetylated layers and completely acetylated layers, respectively. Local disassembly velocity is much lower at completely deacetylated layers as compared to other layers. In addition, the total fraction of lateral layers with *k* ≥ 6 is 0.57, and that with *k* < 6 is 0.43. In contrast to the uniform pattern, these total fractions are not substantially different from each other. Also, the total fraction with *k* ≥ 6 for clustered pattern (0.57) is smaller than that for uniform pattern (0.7). Hence, in filaments with clustered pattern, there are fewer layers which have cooperative increase in local disassembly velocities, compared to the uniform pattern. These factors lead to the smaller values of *mean* disassembly velocities of filaments with clustered patterns observed in Fig 5(c) (blue, black, brown).

The difference in *variance* of disassembly velocities (Fig 5(d)) between uniform and clustered patterns of acetylation can also be understood in terms of the fractions in Fig 7. The higher values of *variance* for the clustered pattern (Fig 5(d), blue) corresponds to the existence of bimodality and a broader distribution in the fraction seen in Fig 7. This arises from the heterogeneity among lateral layers with different values of k, with ample total fraction of layers containing both *k* ≥ 6 and *k* < 6. This results in larger fluctuations in local disassembly velocities. In the case of uniform acetylation pattern, however, there is a single peak in fraction at *k* = 6, with a narrower distribution of fraction over *k*. Hence, disassembly velocities do not have large fluctuations, which results in their *variances* being smaller compared to the clustered patterns (Fig 5(d), green).

## Discussion

Biofilaments such as microtubules are constantly subjected to regulation by Post Translational Modifications (PTM). One of the most widely studied modifications to MTs is acetylation, which is predominantly observed on stable MTs in cells. With a rather slow rate of ≈ 0.03min^−1^, acetylation gradually develops on stable MTs [17–19]. Observations of various patterns of acetylation both *in vivo* and *in vitro* illustrate a scenario in which acetylation proceeds with the tubulin-acetyltransferase enzyme *α*TAT1 entering the MT lumen through open tips or points of defects/lattice openings along the MT filament [15–19]. The ensuing pattern of acetylation may vary depending on the spatial position of entry of *α*TAT1 into the lumen, its binding kinetics and rate of catalysis. *In vitro* experiments have shown that the effect of acetylation is to decrease the lateral interaction between tubulin subunits on adjacent protofilaments on the MT lattice [14]. This means that the existence of various patterns of acetylation is akin to a corresponding distribution of domains of instability across the MT lattice. Cells could thereby utilize relative positions and abundance of acetylation domains on the MT lattice as handles to regulate the state of the microtubule, with each pattern displaying a varying degree of susceptibility to disassembly cues. In this work, we attempt to incorporate the key features of interaction between acetylated tubulin subunits to understand the correlation between acetylation patterns and disassembly profiles of shrinking microtubules.

### Clustered pattern of acetylation ensures smaller *means* and larger local fluctuations of disassembly velocities

In this study, using a multi-protofilament MT lattice, with periodic boundary between the 1^*st*^ and 13^*th*^ protofilaments, we study the effects of three types of acetylation patterns in regulating MT disassembly and the differences between them. The patterns of acetylation we have employed are 1) uniformly random 2) exponentially decreasing from the open tips and 3) clustered at random positions along the MT length. These patterns are observed in experiments under various physiological conditions [15–19]. Results from our simulations suggest that distinct acetylation patterns could result in disassembly profiles that are quite different. For instance, when the filament consists of clustered acetylation patterns along its length, it is more resistant against disassembly compared to a completely uniform or exponentially decreasing pattern of acetylation. When acetylation is distributed completely uniformly across the lattice, it makes the lattice least resistant to disassembly. Further systematics show that this is because in the case of uniform acetylation, acetylated subunits across each layer on the filament act cooperatively to enhance the filament disassembly. In the case of clustered pattern of acetylation, however, there are also several layers which are completely deacetylated and possess lower rates of disassembly. As a result, the filament with clustered acetylation pattern undergoes disassembly with lower values of *mean* velocities (longer *mean* disassembly times) compared to the uniform pattern. The simultaneous existence of large numbers of completely acetylated and completely deacetylated layers in clustered acetylation pattern also leads to higher values of *variance* of disassembly velocities in this case as compared to the uniform pattern.

Theoretical studies have not so far, to the best of our knowledge, investigated the effect of acetylation on MT dynamics within a multi-protofilament model. In this study, we incorporate patterns of acetylation as a heterogeneous disorder distribution of tubulin subunits on the MT lattice. It can be discerned from the results of this study that the inclusion of multiple protofilaments is crucial since the size and spatial distribution of domains of acetylation can regulate the phenomenological parameters associated with filament disassembly.

### Suggested experiments in the context of this study

The distinct patterns of acetylation used in our study are based on observations of the same by *in vitro* experiments under different physiological conditions [15, 18, 19]. Hence, suitable experiments can generate MT filaments with specific patterns of acetylation with the aid of *α*-TAT1 enzymes. In our study, we investigate the disassembly of MT filaments with preformed acetylation patterns, when the free tubulin concentration in the solution is zero. The disassembly dynamics of filaments can be investigated in experiments by diluting the free tubulin concentration to zero (dilution assay), and measuring the times and velocities of shrinkage. However, in order for acetylation patterns to be gradually formed on MTs, these filaments have to be stabilised against disassembly using external proteins while acetylation is in progress. Hence, in order to initiate shrinkage and track the disassembly profiles, the stabilising proteins should also be eliminated from the assay along with free tubulins. From the simulations we obtain differences in mean disassembly velocities upto ≈ 6*μm*/min and variance of disassembly velocities upto ≈ 15*μm*^2^/min^2^, between the uniform pattern and clustered pattern (with *ρ_ld_* = 0.1). These differences are large enough to be measured in dilution assay experiments of MT filaments with distinct preformed acetylation patterns.

Experiments on acetylation have not studied its formation on filaments which undergo polymerisation-depolymerisation kinetics in the absence external stabilising proteins. Although, polymerisation was present in the experiments of Portran *et al* [14], the free tubulin subunits were either completely acetylated or completely deacetylated. While *in vivo* acetylation is predominantly observed on stable MTs, it is also observed to be present to some extent in dynamic MTs as well [44]. The formation of acetylation on a dynamic filament can generate varied patterns of acetylation which can regulate the filament’s shrinkage state. Hence, it is of interest to investigate the coupling of acetylation to the whole polymerisation-depolymerisation kinetics of filaments in future experimental and theoretical studies.

Defects or lattice openings are channels through which the acetyltransferase enzymes may enter the lumen of the filament. Apart from these enzymes, defects are also observed to be the prefered positions at which katanin (a MT severing protein) binds in order to sever the filaments [45]. Moreover, this has been suspected as a mechanism for cells to disassemble the older MTs which have accumulated several defects and also to maintain specific lengths of MTs. Also, *in vivo* experiments have observed a strong preference for katanin to bind to MT filaments with higher levels of acetylation in fibroblasts and dendrites, while not showing the same dependence for axonal MTs [46]. The mean times before severing measured for katanin in Davis *et al* [45] (3.3 ± 2.2min for 180*μ*m filament and 290*nM* katanin; 8.5 ± 4.9min for 73*μ*m filament and 5.7*nM* katanin) are comparable with the maximum difference in disassembly times (≈ 3min) between uniform and clustered acetylation patterns on filaments of ≈ 100*μ*m in our study. Hence, there may be an interesting interplay between the destabilising effects of acetylation and severing proteins on MTs since the distribution of both are affected by the presence of defects. This could be an interesting area of future study.

### Biophysical significance of the study

Transitions between different stages of cell cycle is marked by a distinct shift in MT lengths and numbers. Inadequate disassembly of MT during the prophase has been observed to cause disruption in spindle formation [47]. Various MAPs and severing proteins have been associated with regulating MT stability and disassembly during the various stages of cell cycle.

In this study we have analysed the role of various patterns of acetylation and its total fraction in regulating the stability of filaments against disassembly. It can be discerned from the results that a difference in acetylation patterns can give rise to large differences in MT filament stability, even when the total fraction of acetylation per filament is kept fixed. This becomes more interesting given that under various external conditions, *αTAT*1 turn over rates have been measured to vary by as much as 50 folds [17, 48–50]. An interplay between this and the effective diffusion constant of the enzyme inside the MT lumen will give rise to various patterns of acetylation. Infact, cytoplasmic MTs have been observed to show variability in acetylation patterns. For example, experiments on human fibroblast cells observed that, while a large fraction of cytoplasmic MTs contained acetylated “domains” along the length as well as at the tip, a smaller fraction contained completely uniformly distributed patterns [16]. Results from our study reveal differences in *means* and *variances* of disassembly times and velocities between various acetylation patterns. These differences show that preformed patterns of instabilities on stable lattices can be utilised to regulate response of the lattice to cues to disassemble, as required. Hence, the formation of spatially heterogeneous acetylation patterns can be compared to decisions made by the cell regarding the regulation of filament disassembly before it is triggered.

The state of acetylation of tubulin subunits on MT filaments are observed to affect many cellular structures and functions such as the touch sensation of Touch Receptor Neurons in C.elegans, dynamics of actin filaments, growth of invadopodia, cell migration [48, 51] etc. Kinesin-1 proteins preferentially interact with acetylated tubulins on MTs [52]. Proteins have been observed to preferentially interact with tubulins in other PTM states as well [8, 11]. Based on this, it is proposed that a tubulin “code” may exist, which can be read by proteins associated with MT filaments [53, 54], which can in turn be used for transport or regulation of filament dynamics. However, our results show that, apart from this multifarious effect, characteristic acetylation patterns accumulated under different conditions themselves have signatures which can variably regulate filament disassembly. Hence, in the context of filament disassembly, patterns of acetylation may be a manifestation of the “code” which regulate the phenomenological parameters associated with the MT filament.

## Supporting Information

**S1 text. Determining the parameters used in the simulations**.

In this text we discuss how the parameters used in this work are obtained.

## Acknowledgement

RP acknowledges IYBA, Department of Biotechnology, India for financial support.

## Author Contribution

Conceived and designed the simulations: AJS, RP, DD. Performed the simulations: AJS. Analyzed the data: AJS, RP, DD. Wrote the paper: AJS, RP, DD.

## References

1. Alberts B, Johnson A, Lewis J, Raff M, Roberts K, Walter P. Molecular Biology of the Cell. 4th ed. New York: Garland Science; 2002.

2. Howard J. Mechanics of Motor Proteins and the Cytoskeleton. Massachusetts: Sinauer Associates, Inc.; 2001.

3. Wollman R, Cytrynbaum EN, Jones JT, Meyer T, Scholey JM, Mogilner A. Efficient Chromosome Capture Requires a Bias in the ‘Search-and-Capture’ Process during Mitotic-Spindle Assembly. Curr Biol. 2005;15(9):828 –832. doi:https://doi.org/10.1016/j.cub.2005.03.019.

4. Pearson CG, Yeh E, Gardner M, Odde D, Salmon ED, Bloom K. Stable Kinetochore-Microtubule Attachment Constrains Centromere Positioning in Metaphase. Curr Biol. 2004;14(21):1962 –1967. doi:https://doi.org/10.1016/j.cub.2004.09.086.

5. Sutradhar S, Yadav V, Sridhar S, Sreekumar L, Bhattacharyya D, Ghosh SK, et al. A comprehensive model to predict mitotic division in budding yeasts. Mol Biol Cell. 2015;26(22):3954–3965. doi:10.1091/mbc.E15-04-0236.

6. Mitchison TJ, Kirschner M. Dynamic instability of microtubule growth. Nature. 1984;312:237–242.

7. de Forges H, Bouissou A, Perez F. Interplay between microtubule dynamics and intracellular organization. Int J Biochem Cell Biol. 2012;44(2):266 –274. doi:https://doi.org/10.1016/j.biocel.2011.11.009.

8. Janke C, Chloe Bulinski J. Post-translational regulation of the microtubule cytoskeleton: mechanisms and functions. Nat Rev Mol Cell Biol. 2011;12:773–786. doi:10.1038/nrm3227.

9. Song Y, Brady ST. Post-translational modifications of tubulin: pathways to functional diversity of microtubules. Trends Cell Biol. 2015;25(3):125 –136.

10. Gadadhar S, Dadi H, Bodakuntla S, Schnitzler A, Bièche I, Rusconi F, et al. Tubulin glycylation controls primary cilia length. J Cell Biol. 2017;216(9):2701–2713. doi:10.1083/jcb.201612050.

11. Wloga D, Gaertig J. Post-translational modifications of microtubules. J Cell Sci. 2010;123(20):3447–3455. doi:10.1242/jcs.063727.

12. Sirajuddin M, Rice LM, Vale RD. Regulation of microtubule motors by tubulin isotypes and post-translational modifications. Nat Cell Biol. 2014;16(335).

13. McKenney RJ, Huynh W, Vale RD, Sirajuddin M. Tyrosination of-tubulin controls the initiation of processive dynein–dynactin motility. The EMBO Journal. 2016;35(11):1175–1185. doi:10.15252/embj.201593071.

14. Portran D, Schaedel L, Xu Z, Thery M, Nachury MV. Tubulin acetylation protects long-lived microtubules against mechanical ageing. Nat Cell Biol. 2017;19(4):391 –398.

15. Ly N, Elkhatib N, Bresteau E, Piétrement O, Khaled M, Magiera MM, et al. TAT1 controls longitudinal spreading of acetylation marks from open microtubules extremities. Sci Rep. 2016;6(35624). doi:http://dx.doi.org/10.1038/srep35624.

16. Webster DR, Borisy GG. Microtubules are acetylated in domains that turn over slowly. J Cell Sci. 1989;92(1):57–65.

17. Akella JS, Wloga D, Kim J, Starostina NG, Lyons-Abbott S, Morrissette NS, et al. MEC-17 is an alpha-tubulin acetyltransferase. Nature. 2010;467(7312):218 –222.

18. Agnieszka S, Alexandra, Jeffrey S, Benjamin G, Max, Natasza, et al. Molecular Basis for Age-Dependent Microtubule Acetylation by Tubulin Acetyltransferase. Cell. 2014;157(6):1405 –1415.

19. Coombes C, Yamamoto A, McClellan M, Reid TA, Plooster M, Luxton GWG, et al. Mechanism of microtubule lumen entry for the-tubulin acetyltransferase enzyme TAT1. Proc Natl Acad Sci USA. 2016;113(46):E7176–E7184. doi:10.1073/pnas.1605397113.

20. Cueva J, Hsin J, Huang K, Goodman M. Posttranslational Acetylation of *α*-Tubulin Constrains Protofilament Number in Native Microtubules. Curr Biol. 2012;22(12):1066 –1074. doi:https://doi.org/10.1016/j.cub.2012.05.012.

21. Topalidou I, Keller C, Kalebic N, Nguyen KQ, Somhegyi H, Politi K, et al. Genetically Separable Functions of the MEC-17 Tubulin Acetyltransferase Affect Microtubule Organization. Curr Biol. 2012;22(12):1057 –1065. doi:https://doi.org/10.1016/j.cub.2012.03.066.

22. Howes SC, Alushin GM, Shida T, Nachury MV, Nogales E. Effects of tubulin acetylation and tubulin acetyltransferase binding on microtubule structure. Mol Biol Cell. 2014;25(2):257–266. doi:10.1091/mbc.E13-07-0387.

23. Bowne-Anderson H, Zanic M, Kauer M, Howard J. Microtubule dynamic instability: a new model with coupled GTP hydrolysis and multistep catastrophe. BioEssays. 2013;35(5):452–461. doi:10.1002/bies.201200131.

24. Dogterom M, Leibler S. Physical aspects of the growth and regulation of microtubule structures. Phys Rev Lett. 1993;70:1347–1350. doi:10.1103/PhysRevLett.70.1347.

25. Flyvbjerg H, Holy TE, Leibler S. Microtubule dynamics: Caps, catastrophes, and coupled hydrolysis. Phys Rev E. 1996;54:5538–5560. doi:10.1103/PhysRevE.54.5538.

26. Brun L, Rupp B, Ward JJ, Nedelec F. A theory of microtubule catastrophes and their regulation. Proc Natl Acad Sci USA. 2009;106(50):21173–21178. doi:10.1073/pnas.0910774106.

27. Sumedha, Hagan MF, Chakraborty B. Prolonging assembly through dissociation: A self-assembly paradigm in microtubules. Phys Rev E. 2011;83:051904. doi:10.1103/PhysRevE.83.051904.

28. Padinhateeri R, Kolomeisky AB, Lacoste D. Random hydrolysis controls the dynamic instability of microtubules. Biophys J. 2012;102(6):1274–83. doi:10.1016/j.bpj.2011.12.059.

29. Das D, Das D, Padinhateeri R. Collective force generated by multiple biofilaments can exceed the sum of forces due to individual ones. New J Phys. 2014;16(6):063032.

30. Aparna JS, Padinhateeri R, Das D. Signatures of a macroscopic switching transition for a dynamic microtubule. Sci Rep. 2017;7(45747). doi:https://doi.org/10.1038/srep45747.

31. VanBuren V, Odde DJ, Cassimeris L. Estimates of lateral and longitudinal bond energies within the microtubule lattice. Proc Natl Acad Sci USA. 2002;99(9):6035–6040. doi:10.1073/pnas.092504999.

32. VanBuren V, Cassimeris L, Odde DJ. Mechanochemical Model of Microtubule Structure and Self-Assembly Kinetics. Biophys J. 2005;89(5):2911 –2926. doi:http://dx.doi.org/10.1529/biophysj.105.060913.

33. Martin SR, Schilstra MJ, Bayley PM. Dynamic instability of microtubules: Monte Carlo simulation and application to different types of microtubule lattice. Biophys J. 1993;65(2):578 –596.

34. Molodtsov MI, Ermakova EA, Shnol EE, Grishchuk EL, McIntosh JR, Ataullakhanov FI. A molecular-mechanical model of the microtubule. Biophys J. 2005;88(5):3167–79. doi:10.1529/biophysj.104.051789.

35. Margolin G, Gregoretti IV, Cickovski TM, Li C, Shi W, Alber MS, et al. The mechanisms of microtubule catastrophe and rescue: implications from analysis of a dimer-scale computational model. Mol Biol Cell. 2012;23(4):642–656. doi:10.1091/mbc.E11-08-0688.

36. Jemseena V, Gopalakrishnan M. Microtubule catastrophe from protofilament dynamics. Phys Rev E. 2013;88:032717. doi:10.1103/PhysRevE.88.032717.

37. Zakharov P, Gudimchuk N, Voevodin V, Tikhonravov A, Ataullakhanov FI, Grishchuk EL. Molecular and mechanical causes of microtubule catastrophe and aging. Biophys J. 2015;109(12):2574–2591. doi:10.1016/j.bpj.2015.10.048.

38. Jain I, Inamdar MM, Padinhateeri R. Statistical Mechanics Provides Novel Insights into Microtubule Stability and Mechanism of Shrinkage. PLoS Comput Biol. 2015;11(2):1–23. doi:10.1371/journal.pcbi.1004099.

39. Piedra FA, Kim T, Garza ES, Geyer EA, Burns A, Ye X, et al. GDP-to-GTP exchange on the microtubule end can contribute to the frequency of catastrophe. Mol Biol Cell. 2016;27(22):3515–3525. doi:10.1091/mbc.e16-03-0199.

40. Stukalin EB, Kolomeisky AB. Simple growth models of rigid multifilament biopolymers. J Chem Phys. 2004;121(2).

41. Stukalin EB, Kolomeisky AB. Polymerization dynamics of double-stranded biopolymers: Chemical kinetic approach. J Chem Phys. 2005;122(10):–. doi:http://dx.doi.org/10.1063/1.1858859.

42. Gillespie DT. A general method for numerically simulating the stochastic time evolution of coupled chemical reactions. J Comput Phys. 1976;22(4):403–434. doi:10.1016/0021-9991(76)90041-3.

43. Gardner M, Charlebois B, Jáanosi I, Howard J, Hunt A, Odde D. Rapid Microtubule Self-Assembly Kinetics. Cell. 2011;146(4):582 –592. doi:https://doi.org/10.1016/j.cell.2011.06.053.

44. Perdiz D, Mackeh R, Poüs C, Baillet A. The ins and outs of tubulin acetylation: More than just a post-translational modification? Cell Signal. 2011;23(5):763 –771. doi:https://doi.org/10.1016/j.cellsig.2010.10.014.

45. Davis LJ, Odde DJ, Block SM, Gross SP. The Importance of Lattice Defects in Katanin-Mediated Microtubule Severing in Vitro. Biophys J. 2002;82(6):2916 –2927. doi:https://doi.org/10.1016/S0006-3495(02)75632-4.

46. Sudo H, Baas PW. Acetylation of Microtubules Influences Their Sensitivity to Severing by Katanin in Neurons and Fibroblasts. J Neurosci. 2010;30(21):7215–7226. doi:10.1523/JNEUROSCI.0048-10.2010.

47. Mchedlishvili N, Matthews HK, Corrigan A, Baum B. Two-step interphase microtubule disassembly aids spindle morphogenesis. BMC Biology. 2018;16(1):14. doi:10.1186/s12915-017-0478-z.

48. Shida T, Cueva JG, Xu Z, Goodman MB, Nachury MV. The major-tubulin K40 acetyltransferase *α*-TAT1 promotes rapid ciliogenesis and efficient mechanosensation. Proc Natl Acad Sci USA. 2010;107(50):21517–21522. doi:10.1073/pnas.1013728107.

49. Friedmann DR, Aguilar A, Fan J, Nachury MV, Marmorstein R. Structure of the *α*-tubulin acetyltransferase, TAT1, and implications for tubulin-specific acetylation. Proc Natl Acad Sci USA. 2012;109(48):19655–19660. doi:10.1073/pnas.1209357109.

50. Davenport AM, Collins LN, Chiu H, Minor PJ, Sternberg PW, Hoelz A. Structural and Functional Characterization of the *α*-Tubulin Acetyltransferase MEC-17. J Mol Biol. 2014;426(14):2605 –2616. doi:https://doi.org/10.1016/j.jmb.2014.05.009.

51. Mirvis M, Stearns T, James Nelson W. Cilium structure, assembly, and disassembly regulated by the cytoskeleton. Biochem J. 2018;475(14):2329–2353. doi:10.1042/BCJ20170453.

52. Reed NA, Cai D, Blasius TL, Jih GT, Meyhofer E, Gaertig J, et al. Microtubule Acetylation Promotes Kinesin-1 Binding and Transport. Curr Biol. 2006;16(21):2166 –2172. doi:https://doi.org/10.1016/j.cub.2006.09.014.

53. Janke C, Kneussel M. Tubulin post-translational modifications: encoding functions on the neuronal microtubule cytoskeleton. Trends Neurosci. 2010;33(8):362 –372. doi:https://doi.org/10.1016/j.tins.2010.05.001.

54. Janke C. The tubulin code: Molecular components, readout mechanisms, and functions. J Cell Biol. 2014;206(4):461–472. doi:10.1083/jcb.201406055.

